# Developing a network view of type 2 diabetes risk pathways through integration of genetic, genomic and functional data

**DOI:** 10.1101/350181

**Authors:** Juan Fernéndez-Tajes, Kyle J Gaulton, Martijn van de Bunt, Jason Torres, Matthias Thurner, Anubha Mahajan, Anna L Gloyn, Kasper Lage, Mark I McCarthy

**Author notes:** These authors equally contributed to this work (Juan Fernéndez-Tajes, Kyle J Gaulton). Corresponding author: Mark I McCarthy;. Department of Bioinformatics and Data Mining, Novo Nordisk A/S, Maaloev, Denmark (Kyle J Gaulton, Martijn van de Bunt).

## Abstract

Genome wide association studies (GWAS) have identified several hundred susceptibility loci for Type 2 Diabetes (T2D). One critical, but unresolved, issue concerns the extent to which the mechanisms through which these diverse signals influencing T2D predisposition converge on a limited set of biological processes. However, the causal variants identified by GWAS mostly fall into non-coding sequence, complicating the task of defining the effector transcripts through which they operate. Here, we describe implementation of an analytical pipeline to address this question. First, we integrate multiple sources of genetic, genomic, and biological data to assign positional candidacy scores to the genes that map to T2D GWAS signals. Second, we introduce genes with high scores as seeds within a network optimization algorithm (the asymmetric prize-collecting Steiner Tree approach) which uses external, experimentally-confirmed protein-protein interaction (PPI) data to generate high confidence subnetworks. Third, we use GWAS data to test the T2D-association enrichment of the “non-seed” proteins introduced into the network, as a measure of the overall functional connectivity of the network. We find: (a) non-seed proteins in the T2D protein-interaction network so generated (comprising 705 nodes) are enriched for association to T2D (*p*=0.0014) but not control traits; (b) stronger T2D-enrichment for islets than other tissues when we use RNA expression data to generate tissue-specific PPI networks; and (c) enhanced enrichment (*p*=3.9×l0^−5^) when we combine analysis of the islet-specific PPI network with a focus on the subset of T2D GWAS loci which act through defective insulin secretion. These analyses reveal a pattern of non-random functional connectivity between causal candidate genes atT2D GWAS loci, and highlight the products of genes including *YWHAG*, *SMAD4* or *CDK2* as contributors to T2D-relevant islet dysfunction. The approach we describe can be applied to other complex genetic and genomic data sets, facilitating integration of diverse data types into disease-associated networks.

**Author summary:** We were interested in the following question: as we discover more and more genetic variants associated with a complex disease, such as type 2 diabetes, will the biological pathways implicated by those variants proliferate, or will the biology converge onto a more limited set of aetiological processes? To address this, we first took the 1895 genes that map to ~100 type 2 diabetes association signals, and pruned these to a set of 451 for which combined genetic, genomic and biological evidence assigned the strongest candidacy with respect to type 2 diabetes pathogenesis. We then sought to maximally connect these genes within a curated protein-protein interaction network. We found that proteins brought into the resulting diabetes interaction network were themselves enriched for diabetes association signals as compared to appropriate control proteins. Furthermore, when we used tissue-specific RNA abundance data to filter the generic protein-protein network, we found that the enrichment for type 2 diabetes association signals was enhanced within a network filtered for pancreatic islet expression, particularly when we selected the subset of diabetes association signals acting through reduced insulin secretion. Our data demonstrate convergence of the biological processes involved in type 2 diabetes pathogenesis and highlight novel contributors.

## Introduction

The rising prevalence of type 2 diabetes (T2D) represents a major challenge to global health [1]. Current strategies for both prevention and treatment of T2D are suboptimal, and greater insight into the mechanisms responsible for the development of this condition is a prerequisite for further advances in disease management [2].

The identification of human DNA sequence variants which influence predisposition to T2D provides one of the most direct approaches for deriving mechanistic insight. However, current understanding of the genetic architecture of T2D indicates that the genetic component of T2D predisposition likely involves variation across many thousands of loci [3, 4], Close to 500 independent genetic signals for which there is robust evidence of a contribution to T2D predisposition have been identified, largely through genome-wide association studies, supplemented by analysis of exome- and genome-sequence data [4–6]. This profusion of genetic signals has raised questions concerning the extent to which the inherited susceptibility to complex traits such as T2D can be considered to occupy finite biological space [7], In other words, as the number of loci influencing T2D risk increases, will the mechanisms through which these are found to mediate the development of this condition continue to proliferate, or will they start to converge around a limited set of pathways?

There are two main challenges in addressing this key question. First, whilst a minority of the causal variants underlying these association signals are coding (and therefore provide direct inference regarding the genes and proteins through which they act), most lie in regulatory sequence. This makes assignment of their effector transcripts a non-trivial exercise, and obscures the downstream mechanisms through which these variants impact T2D-risk [8–10]. This challenge can increasingly be addressed through the integration of diverse sources of relevant data including (a) experimental data (e.g. from studies of cis-expression or conformational capture) which link regulatory risk-variants to their likely effectors [11,12]; and (b) evaluations of the biological evidence connecting each of the genes within a GWAS-associated region to the disease of interest. In the present study, focussing on a set of approximately 100 T2D-risk loci with the largest effects on T2D predisposition, we use a range of information to derive “positional candidacy” scores for each of the coding genes mapping to T2D-associated GWAS intervals.

The second challenge lies in the requirement to define functional relationships between sets of candidate effector transcripts in ways that are robust, and, in particular, orthogonal to the data used to assign candidacy in the first place [13,14], Solutions for the second challenge are less well-developed but generally involve some type of network analysis (e.g. weighted gene correlation network analysis [WGCNA]) and application of the “guilt-by-association” framework to infer function [15–17], However, recourse to co-expression information, or functional pathway enrichment methods to generate and evaluate such networks runs the risk of introducing circularity, given that information on expression and function typically contributes (whether explicitly or not) to assignments of effector transcript candidacy. The use of protein-protein interaction data provides one possible solution to this conundrum [18]. In the present study, we make use of external protein-protein interaction data from the lnWeb3 dataset [19, 20] to evaluate and characterise the connectivity of the T2D candidate effector transcripts in terms of their ability to nucleate empirically-confirmed interactions between their encoded proteins.

## Materials and methods

### Positional candidacy score derivation

We developed a framework to score the candidacy of genes mapping to GWAS association signals which aggregated data from multiple sources. The information collected fell into two categories. First, we used regression-based approaches to link disease-associated variants (most of which map into non-coding sequence and are therefore presumed to act through transcriptional regulation of nearby genes) to their likely effector transcripts, using a combination of variant-based annotations and expression QTL data. Second, we scored each of the genes in these GWAS regions for disease-relevant biological function. We combined the two measures to generate a “positional candidacy score” (PCS) for each gene. We applied this framework to 1895 genes located within a 1Mb interval around the lead variants from 101 T2D GWAS regions. These represent the loci with the largest effect sizes for T2D, as identified in European subjects as of early 2017 [4, 6, 21]: Supplementary Table 1). The 1Mb intervals contained 1895 genes.

### Mapping effector transcripts to GWAS signals

At each of the 101 loci, we collected summary T2D case-control association data (−log_10_p values) for all 1000 Genomes variants in the 1Mb interval surrounding the lead variants [6]. We then annotated variants in each interval using gene-based annotations for all genes in the interval from several sources. First, we collected relevant discrete annotations for all protein coding genes in GENCODE (version 19) [22] within the interval including (a) coding exon location; (b) promoter location (defined as lkb region upstream of the transcription start site [TSS]); (c) distal regulatory elements correlated with gene activity from DNAsel hypersensitivity (DHS) data (ENCODE version 3) [23]. We assigned each variant a binary value based on whether it overlapped one of the discrete annotations for a gene in the interval (exon, promoter, distal element). Second, we collected summary statistic expression QTL (eQTL) data from liver, skeletal muscle, whole blood, subcutaneous adipose and visceral adipose (GTEx version 6) [24] and pancreatic islets [11]. We assigned each variant the −log_10_p value of eQTL association for each cell type for each gene in the interval. Third, we calculated the distance of each variant to the TSS of each gene in the interval, and assigned each variant the inverse TSS distance for each gene (i.e. variants closer to the TSS have higher values). Variants without values in the eQTL datasets were removed from the analysis.

We then performed feature selection for each T2D locus separately using elastic net regression (R package glmnet) with the T2D p-values as the outcome variable and binary genomic annotations (exon, promoter, distal element), distance to TSS, and cell type cis-eQTL p-values for each gene in the interval as the predictor variables. We also included minor allele frequency and imputation quality of each variant at the locus as predictor variables. We obtained the effects of features selected from the resulting model. We applied a 10-fold scaling factor to coding exon features, based on known enrichment of T2D variants in coding exons [25, 26]. Where multiple features were selected for the same gene (e.g. distal DHS site and tissue eQTL) we summed the effects for that gene. We considered the summed effects of features for each gene as the ‘variant link score’ in subsequent analyses.

### Semantic mapping of gene functional annotations

We also derived a second score of the T2D-relevance for each gene within the 101 GWAS intervals based on the annotations for each within data from gene ontology (GOA, version 157), the mouse genome database (MGD, version 6.08), and biological pathways (KEGG) (version 83.1), compiling these annotations into a single document per gene. We also created a query document of empirically-compiled terms we considered relevant to T2D pathophysiology (listed here: https://github.com/kjgaulton/gene-pred/blob/master/res/T2D.query.manual.txt). Both gene documents were converted into a word matrix. We calculated the total number of unique words *a*cross all documents *N*, after removing a list of commonly used “stop” words from PubMed (https://www.ncbi.nlm.nih.gov/pubmed/) and stemming the remaining words. We weighted each word *w* for each gene document *g* using “term frequency (TF)” minus “inverse document frequency” defined as:

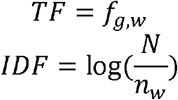

where *n*_*w*_ is the number of documents containing word *w*. We defined the value (*g*_*w*_) of word *w* in gene document *g* as:

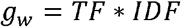

and applied latent semantic analysis (LSA) using singular value decomposition of the weighted matrix *M*

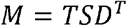

where *T* is the left singular vector matrix of terms, *D* is the right singular vector matrix of documents, *S* is the diagonal matrix of singular values, and the number of dimensions was determined by the function *dimcalc_share* from the *lsa* package [27]. We used the resulting matrices to identify genes with functional attributes that indicated relevance to T2D pathogenesis. For each gene document vector *g*, we calculated similarity scores S_*i*,*q*_ using the dot product between the gene vector and the T2D query vector *q*

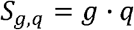

From these data, we extracted similarity scores for the 1895 genes of interest, which we considered the ‘semantic score’ in subsequent analyses.

### Combining gene scores

For each of the 1895 genes, we scaled scores from these two analyses to the sum of scores for each of the *x* genes at each locus resulting in a semantic score *s*_*g*_ and variant link score *v*_*g*_. To calculate a positional candidacy score (PCS), we averaged the two scores and rescaled across all *x* genes at each locus.

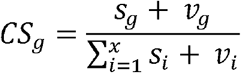

### Network modelling

#### Selection of the “seed node set”

At each GWAS locus, we defined the sets of genes that, after ranking the genes for each locus by decreasing PCS, generated a cumulative PCS exceeding 70%. This reduced the set of 1895 genes of interest to 451 “seed” nodes for subsequent network analysis. We performed network analyses using an updated version of InWeb3, a previously-described comprehensive map of protein-protein interactions, containing 169,736 high-confidence interactions between 12,687 gene products compiled from a variety of sources [19, 20]. We updated the version used in [20], by updating outdated gene symbols and restricting interactions to those deemed “high-confidence” (score >0.124).

#### Prize-collecting Steiner Tree formulation

We formulated the task of examining the connectivity of GWAS positional candidates (the set of 451 “seed” genes) within protein-protein interaction space as an asymmetric prize-collecting Steiner tree (APCST) problem. APCST-like approaches have been widely used to solve network-design problems [28–30]. The APCST seeks to connect “seed” nodes (in formal nomenclature, “terminals”) to collect “prizes”, using confirmed protein-protein interactions as edges. Prizes are weights added to seed nodes: in our analysis, these correspond to the PCS values for each “seed” gene, derived from the -omic integration approach. “Linking” (formally, “Steiner”) nodes (that is, proteins/genes not included in the seed set) can be introduced into the network, where necessary. Network expansion is controlled by the balance between the benefits of adding a particular node (increased connectivity between seed genes, driven by the collection of prizes) vs. the costs of adding additional edges (based on a function which penalises expansion of the network). In mathematical terms, we defined the APCST as follows: given a directed graph *G* = (*V, A*), arc costs *c*: *A* ⟼ ℝ ≥ 0, node prizes *p*: *V* ⟼ ℝ ≥ 0 and a set of fixed terminals *T*_*f*_ the goal is to find an arborescence *S* = (*V*_*s*_,*A*_*s*_) ⊆ *G* that spans *T*_*f*_ such that the following function is maximized:

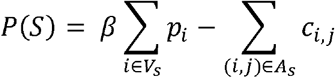

In this formulation, we reward the inclusion of nodes *i* ∈ *V*_*s*_ with higher prizes (that is, higher PCS values) (first term of equation) while paying costs for including edges (second term of equation). The parameter, *β*, scales the importance of node prizes versus edge costs in the optimization and can be used to titrate the size of the generated network. We tested different values of *β* (between 4 and 30) and selected *β* =8 that produced a manageable network size (~130 genes) and included > 25% of the seed node set (**SI Fig**).

Although this problem is NP-hard (nondeterministic polynomial time-hard) [31], the APCST algorithm is found to be efficient in calculating exact and proximal solutions (DIMACs 11^th^ challenge, http://dimacsll.zib.de/). The branch-and-bound algorithm, implemented in *dapstp* algorithm (https://github.com/mluipersbeck/dapcstp) and using default parameters, was used to find the optimal (or near optimal) APCST solution.

### Generation of networks using dapcst algorithm

We used a particular variation of the ACPST (“root-ACPSTP”) where the search for the optimal solution starts in a specific node. This allowed us to force each seed node in turn to be included in the network, in contrast to the default APCST method which initialises network construction from the nodes with higher weights. For the main T2D analysis, therefore, the algorithm was run 451 times, once for each “seed” node. Runs generating a network of >10 nodes (353 networks, median 155 nodes) were combined to form an ensemble network from the union of all *n* networks. This was reprojected onto the lnWeb3 interactome to recover missing connections across nodes. As this final network represents a superposition of many different networks, linking nodes may sometimes appear at the periphery.

We assessed the specificity of each node in the final network by running the algorithm 100 times with the same parameter settings, but with random input data. We define specificity in this context as the complement of the percentage with which a given seed or linking node from the final network appears in runs generated from random input data. For each random run, we selected, from the lnWeb3 interactome, random seed nodes matching the binding degree distribution of the observed set of seeds, and assigned them the same prize value as the original. Using the final parameter settings, we found that the included linking nodes were highly specific to our particular data, with 80% of them having a specificity higher than 75% (**S2 Fig**).

### Testing network for Enrichment in GWAS signal

To evaluate the extent to which the PPI network provided functional connectivity between positional candidates across loci, we measured the enrichment of the linking nodes forT2D association signals. This avoided the circularity of using co-expression or functional data to evaluate connectivity (as both contributed to the PCS determination). We generated gene-wise p values using the PASCAL method [32] from large-scale GWAS studies across a set of 33 traits (using data extracted from public repositories) including a recent meta-analysis of T2D GWAS data from ~150,000 Europeans [6]. We mapped these gene-wise association p-values to linking nodes, and converted them to Z-scores using the standard normal cumulative distribution, *Z*_i_ = *ϕ*^−1^(1 − *p*_*i*_). We then quantified GWAS enrichment by aggregating the Z-scores using Stouffer’s method:

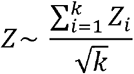

where *Z*_*i*_ is the Z-score for the gene-wise p value for linking node *i* and *k* is the number of linking nodes in the network. Then, by permuting the InWeb3 network using a node permutation scheme, we compared the observed enrichment in GWAS signals to a random expectation, allowing us to calculate a nominal p value as:

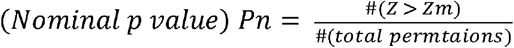

where *p*_*n*_ is the permuted p value generated in the permutation scheme. In this last step, the binding degree of all genes in the network is taken into full consideration (i.e. they all have the same binding degree as provided by the APCST network). To minimise bias arising from the co-localisation of genes with related functions (which is a feature of some parts of the genome), in each of these permutations we only considered proteins whose genes mapped outside a 1Mb window around the lead SNP for any significant GWAS association for that trait.

### APCST model clustering

To aid interpretation of the PPI networks, we used a community clustering algorithm that maximizes network modularity and which breaks the full APCST model into smaller subnetworks [33].

### GTEx and Islet RNAseq datasets

The InWeb3 PPI network we used is generated from empirically-confirmed interactions, but nevertheless includes many interactions that, owing to restricted tissue-specific expression, are unlikely to be biologically relevant. We used tissue-specific RNA expression data to filter the overall lnWeb3 network and thereby generate *in silico* “tissue-specific” PPI networks, using TPM counts from GTEx (version 7: https://www.gtexportal.org/home/, last accessed 21 Oct 2017), complemented by human pancreatic islet data from [11], Proteins with mRNA TPM counts <1 in over 50% of samples for that tissue were removed from the InWeb network, allowing us to generate in silico PPI networks for 46 tissues.

### Functional Enrichment Analysis

Gene Set Enrichment (GSE) of networks and sub-networks were assessed with ClueGO [34] using GO terms and REACTOME gene sets [35]. The enrichment results were grouped using a Cohen’s Kappa score of 0.4 and terms were considered significant when Bonferroni adjusted p-value <0.05 and at least 3% of the genes contained in the tested gene set were included in the network. Cohen’s Kappa statistic measures the gene-set similarity of GO terms and REACTOME pathways and allowed us to group enriched terms into functional groups that improve visualization of enriched pathways.

## Results

### Prioritizing positional candidates at T2D risk loci

We implemented a framework to derive positional candidacy scores (PCS) for genes within T2D GWAS loci through the aggregation of two main types of data (**Fig. 1**; **Methods**). First, we used regression-based approaches to link disease-associated variants (most of which map to non-coding sequence and are therefore presumed to act through transcriptional regulation of nearby genes) to their likely effector transcripts, using a combination of variant-based annotations and expression QTL data. Second, we scored each of the genes in these GWAS regions for disease-relevant biological function using semantic mapping of gene functional annotations from Gene Ontology, Mouse Genome Database, and KEGG. We combined the evidence from both approaches, normalized across all genes at each GWAS locus, to generate the PCS for each gene.

**Figure 1.**
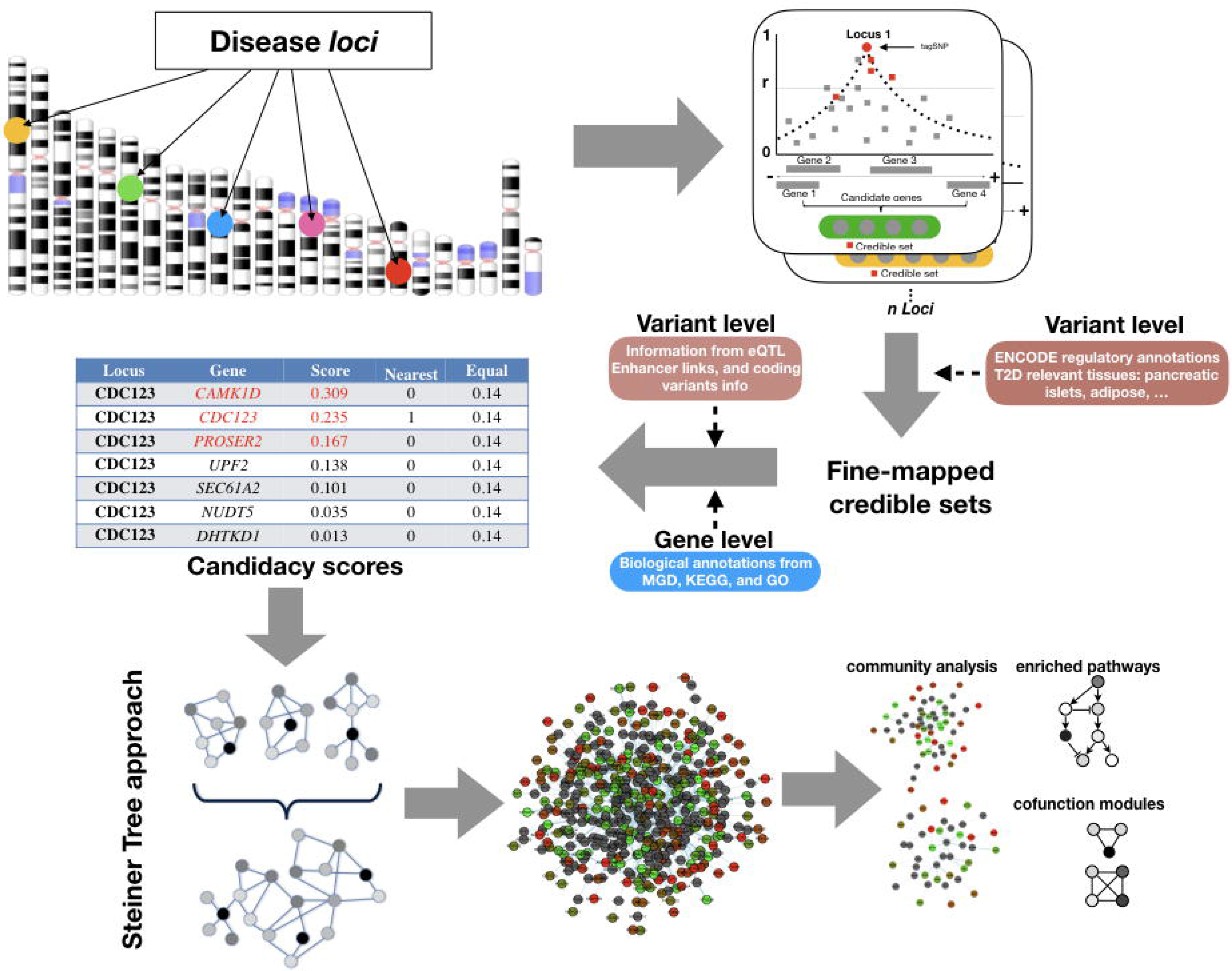
Overview of the Data Integration pipeline. We collected variants in the 1Mb interval surrounding index variants at each of the 101 T2D GWAS loci along with relevant annotations for all protein coding genes in GENCODE including coding exon location, promoter location, distal regulatory elements correlated with gene activity from DNAsel hypersensitivity (DHS) data and summary statistic expression QTL (eQTL) data from T2D-relevant tissues. This, combined with information at gene level from a semantic similarity metric, allowed us to define positional candidacy scores for each gene in the GWAS intervals. These genes were projected into the lnWeb3 data set using a Steiner tree algorithm to define a PPI network that maximises candidate gene connectivity. This network was further analysed to find processes, pathways, and genes implicated in T2D pathogenesis.

We applied this method to score 1,895 genes mapping within a 1Mb interval around the lead variant at 101 T2D GWAS regions. This list of 101 T2D loci was assembled from a series of recent large-scale T2D GWAS studies [4, 6, 21] and represents the largest-effect T2D GWAS loci identified as of early 2017. The 1Mb interval was selected to capture the majority of cis-acting regulatory effects (95% of cis-eQTLs map within 445kb of the lead SNP [24]), and is therefore also likely to encompass most potential effector genes [36]. We observed only weak correlation between the semantic and risk variant link scores for the 1,895 positional candidates (r^2^=0.05, p=0.01), indicating that these provide distinct information (**S3 Fig**).

Most (71%) of the 1,895 genes had minimal evidence linking them to a causal role in T2D pathogenesis (PCS<0.05) (**S3 Fig**). However, 95% of T2D loci included at least one gene (median, 3) with PCS>0.10, and at 70% of loci, there was at least one gene with PCS>0.20 (**S3 Fig**). The top-scoring genes across the 101 loci (such as *IRS1* [PCS=0.69], *SLC30A8* [PCS=0.77], *HNF1B* [PCS=0.54]) include many of the genes with the strongest prior claims for involvement in T2D risk, prior claims which arise in part from data used to generate the PCSs. For example, these genes each contain rare coding variants directly implicated in development of T2D (or related conditions): these variants are independent of the common variant GWAS signals, but their relationship to diabetes is likely to have been captured through the semantic mapping. The PCS also highlighted several other highly-scoring candidates with known causal roles in relation to diabetes and obesity such as *MC4R* (PCS=0.43), *WFS1* (0.41), *ABCC8* (0.37), *LEP* (0.27), *GCK* (0.24), and *HNF1A* (0.23). At other loci, these analyses highlighted candidates that have received scant attention to date: for example, *CENPW* (PCS=0.83) scored highly both in terms of semantic links to T2D-relevant processes and an adipose cis-eQTL linking the T2D GWAS SNP to *CENPW* expression.

To define the seed-genes for subsequent PPI analyses, we gathered the sets of genes that, after ranking the transcripts for each locus by decreasing PCS, cumulatively accounted for at least 70% of the candidacy score for each locus. For example, at the *TP53INP1* locus, where the gene-specific PCSs range from 0.01 to 0.16 across a total of 17 mapped genes, the seed-gene set includes the first six (**S4 Fig**). This filter identified a total of 451 positional candidates across the loci, reducing the median number of genes per locus from 19 to 6 (**S4 Fig**). This filtering mostly removes genes with low PCS values: the proportion of genes with PCS<0.05 falls from 71% to 12%, while most genes with PCS>0.1 or >0.2 are retained (**S3 Fig**).

This prioritisation process ensures that genes with the strongest combined causal evidence are favoured for network modelling, resulting in sets of seed genes that are more extensive than selection based on proximity alone (such as “nearest gene” approaches that seek to generate networks from only the genes mapping closest to the lead variants) but smaller than those which consider all regional genes of equal weight (“all gene” approaches). Note that our strategy does not require complete ascertainment of all true causal genes within this set of 451 genes: true effector genes excluded from the prioritised set of 451 genes (e.g. because they map more distal to the lead variant than 500kb) remain available for “discovery” through the network modelling described below.

### Building a T2D-relevant protein-protein interactome

We set out to test whether this list of prioritised candidates could be used to characterise the functional relationships between genes (and proteins) implicated in T2D pathogenesis. Because the PCS scores used to prioritise the genes already incorporated (explicitly or otherwise) diverse types of functional and expression data, biasing any assessment of connectivity in these domains, we focused the network analysis around protein-protein interaction (PPI) data. To do so, we projected these 451 genes onto externally-derived, empirically-driven PPI resources (lnWeb3) [19, 20] using an established network modelling strategy (the Asymmetric Prize-Collecting Steiner Tree (APCST)) (**Fig1**; **Methods**). In this analysis, the 451 positional candidates represent “seed” nodes which are used by the APCST algorithm to generate PPI networks which seek (with appropriate penalties to prevent frivolous propagation) to connect as many seed nodes as possible to each other, either directly, or using other (non-seed) proteins as links (“linking” nodes). The network topology is dependent only on the PCS values of the “seed” genes which are carried forward as weights into the APCST analysis, the confidence scores for each of the empirical PPI interactions in lnWeb3, and the beta value used to tune the overall size of the PPI network generated (see **Methods)**.

We operationalised the PPI network as follows (see **Methods**). Using each “seed” gene in turn, we used InWeb3 data to generate a PPI network that maximised the connectivity to other seed genes within the constraints of the APCST model. Of the 451 seed genes, 98 failed to produce a network exceeding 10 nodes. The remaining 353 networks had a median of 110 seed and 45 linking nodes and were combined into an ensemble network, which was again projected into the InWeb3 interactome to recover missing connections between nodes. The final network contained 705 nodes (431 seed nodes, 274 linking nodes) and 2678 interactions (**Fig 2**). Based on random networks generated with the same algorithm (see **Methods**), 80% of the linking nodes have a specificity for membership of the final network exceeding 75%, indicating that these linking nodes do not simply reflect generic hubs in PPI space (**S2 Fig**).

**Figure 2.**
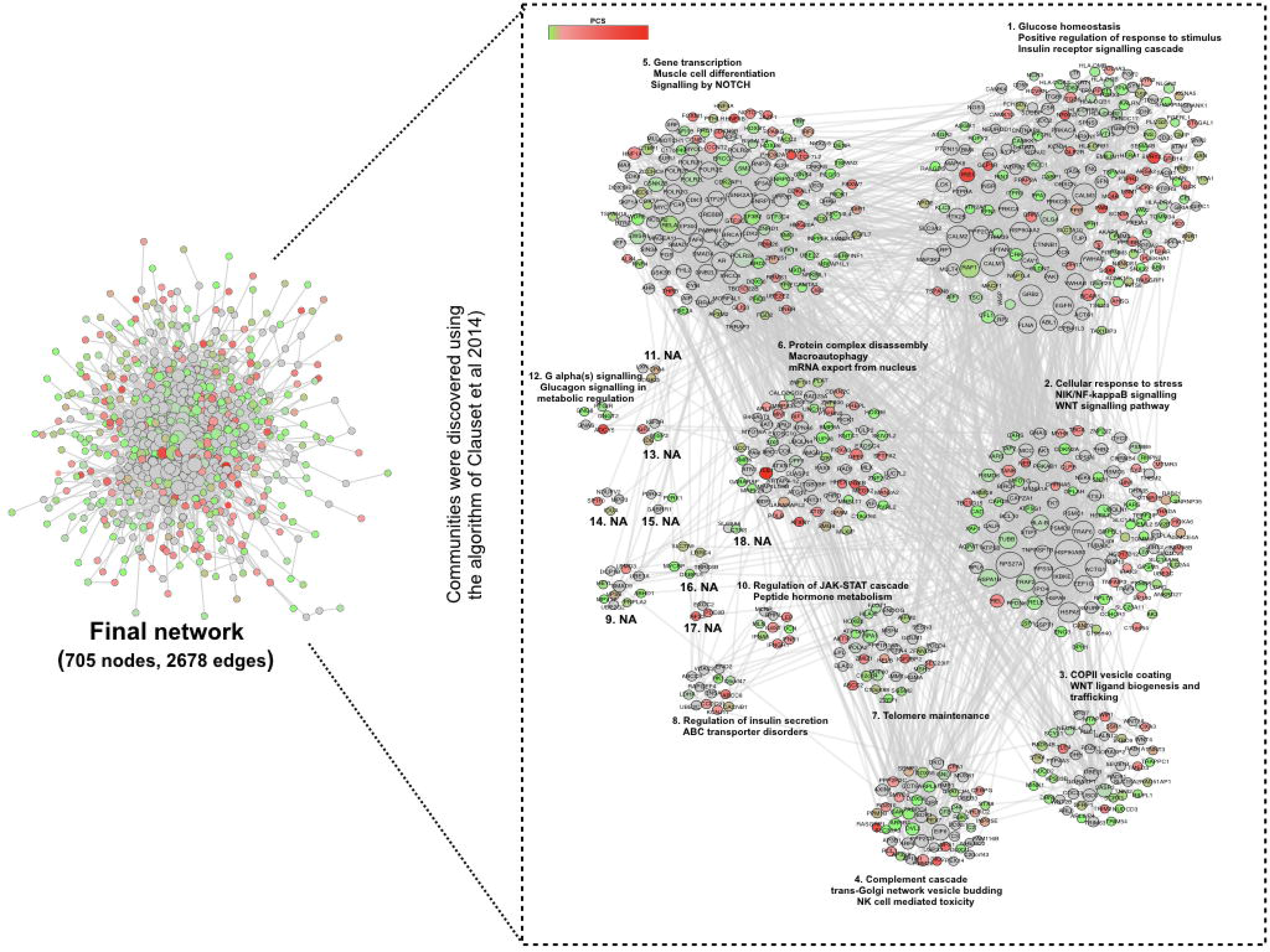
APCST final network. The final PPI network generated from the T2D GWAS interval genes includes 431 seed nodes and 274 linking nodes connected by 2,678 interactions. We divided this network into 20 sub-networks (communities) using a community clustering algorithm that maximizes network modularity (33), and highlighted enrichment of specific biological processes contained within these based on Gene Ontology terms and REACTOME pathways. Coloured nodes represent seed nodes, whereas grey nodes represent linking nodes.

### The T2D PPI network is enriched for T2D associations

If the final network truly provides novel insights into the functional relationships between genes thought to be mediating T2D predisposition, we reasoned that the “linking” genes (those brought into the network purely on the basis of external data indicating their protein-level interaction with seed genes) should be enriched for other seed gene characteristics. To avoid circularity arising from validation using data types that had contributed to the generation of the original PCS weights, including measures of gene function (eg GO, KEGG) or RNA expression data, we turned to T2D GWAS data, looking for evidence that the genes encoding the linking proteins were themselves enriched for T2D-association signals. For this, we used T2D-association data from a set of ~150,000 European T2D case-control subjects imputed to 1000 Genomes [6]. Briefly, the linking nodes were mapped to gene-wise association p-values generated from the GWAS results using PASCAL [32], The significance of the collective enrichment of these gene-wise p-values was obtained by permuting the observed set of linking nodes with equivalent sets of “random” nodes from the lnWeb3 database, matched for binding degrees (see **Methods**). To minimise the prospects of picking up false signals arising from the combination of local LD and the nonrandom genomic location of functionally-related genes, we excluded all genes from the IMb-window around the 101 lead variants from these analyses.

Compared to the distribution of scores in the permuted background, the gene-wise p-values for linking genes in the empirical reconstructed network demonstrated significant enrichment of T2D association (p=0.0014). To confirm that this enrichment was specific to T2D, we repeated the analysis, retaining the same PPI final network, but instead using GWAS data (and PASCAL-derived gene-wise p-values) from 33 different traits across a wide range of disease areas. The only other traits displaying evidence of GWAS enrichment within the linking nodes of the T2D PPI network were those for anthropometric traits with known relevance to T2D pathophysiology (**S5 Fig**).

To gain insights into how the linking nodes of our final network contribute to T2D biology, we used the DisGeNET database [37], which collates gene-disease information from public data as well as from literature via natural language processing tools. We focused on the 274 linking nodes included in our model to avoid circularity arising from using the seeds, and identified 92 (~33%) with known links to T2D (**S2 Table**). Examples include: (a) *NEUROD1* which encodes a transcription factor that is involved in the development of the endocrine cell lineage and has been implicated in monogenic diabetes [38]; (b) *PRKCB* involved in insulin resistance [39], and (c) *GNAS*, implicated in beta-cell proliferation [40]. For this last gene, mice knockouts have been shown to produce phenotypes concordant with diabetes [41]. These examples demonstrate the potential of these analyses to draw in “linking” nodes as related to T2D even when they are not located within genome-wide association signals.

### The T2D PPI network captures biological processes relevant to disease pathogenesis

To increase biological interpretability, we next sought to split the large final PPI network of 705 nodes into smaller sub-networks of closely-interacting proteins (“communities”). Using the algorithm proposed by [33], we identified 18 such communities (each containing between 2 and 186 nodes) (**Fig 2**). We performed enrichment analyses on each community using GO and REACTOME datasets, this time including both seed and linking nodes. We observed that the individual sub-networks were enriched for processes including “glucose homeostasis” and “insulin receptor signalling cascade” (sub-network 1), “Wnt” and “NIK/NF-kappaB signalling pathways” and “cellular response to stress” (subnetwork 2), “COPII vesicle coating” and “Wnt ligand biogenesis and trafficking” (sub-network 3), “regulation of insulin secretion” (sub-network 8), and “glucagon signalling in metabolic regulation” (sub-network 12) (**Fig 2**, **S3 Table**). This pattern of functional enrichment is broadly consistent with existing knowledge regarding aspects of T2D pathogenesis [42–44], We saw no evidence in support of certain processes that have been proposed as contributors to T2D pathogenesis such as mitochondrial function, or oxidative phosphorylation [45, 46], in line with the paucity of evidence linking these processes to T2D risk in standard gene-set enrichment analyses [4, 21].

### Information on tissue-specificity enhances the model

The APCST model described above was constructed from a generic, tissue-agnostic PPI network. As a result, it features edges that, whilst they may be supported by the empirical data used to generate the InWeb3 database, are unlikely to be pathophysiologically relevant, due to mutually-exclusive tissue-specific expression patterns. We hypothesised that the use of tissue-specific interactomes, focused on T2D-relevant tissues, would allow us to refine the reconstructed PPI network, and might enhance the GWAS enrichment signal. In the absence of empirical PPI data for all relevant tissues, we generated these tissue-specific PPI networks by filtering on RN A transcript abundance. Starting from the generic final APCST network, we removed, for each tissue, all nodes (and their corresponding edges) with little or no transcriptional activity (see **Methods)**. In all, we generated tissue-specific PPI networks, using RNA-Seq data sourced from 46 different tissues, 45 (including fat, liver and skeletal muscle) from GTEx (v7) [24][www.gtexportal.org] (median number of individuals = 235) together with a set of human islet RNA-seq data (n=118) [11], which had been reprocessed through a GTEx-aligned pipeline.

We then repeated the T2D GWAS signal enrichment analysis (“linking” nodes only; 100,000 permutations) across each of these 46 tissue-specific PPI networks. We detected broad enrichment for T2D association in linking nodes across many of these tissue-specific networks: this likely reflects the fact that these tissue-specific networks remain highly overlapping (**S6 Fig**). Nonetheless, with the exception of whole blood, the strongest enrichment signal for T2D GWAS data was observed in the islet-specific PPI network (**Fig 3**). This enrichment was less significant (p=0.019) than that observed in the full network (p=0.0014), but this, at least in part, reflects the reduction in the number of linking nodes in the islet-specific network (from 274 to 229). Other tissues implicated in T2D pathogenesis such as adipose, skeletal muscle or liver generated more limited evidence of enrichment (**Fig 3**). This pattern of enrichment (favoring islets, and to a lesser degree, adipose) mirrors equivalent observations for other tissue-specific annotations (including cis-eQTL signals and active enhancers) with respect to T2D association data [10,11].

**Figure 3.**
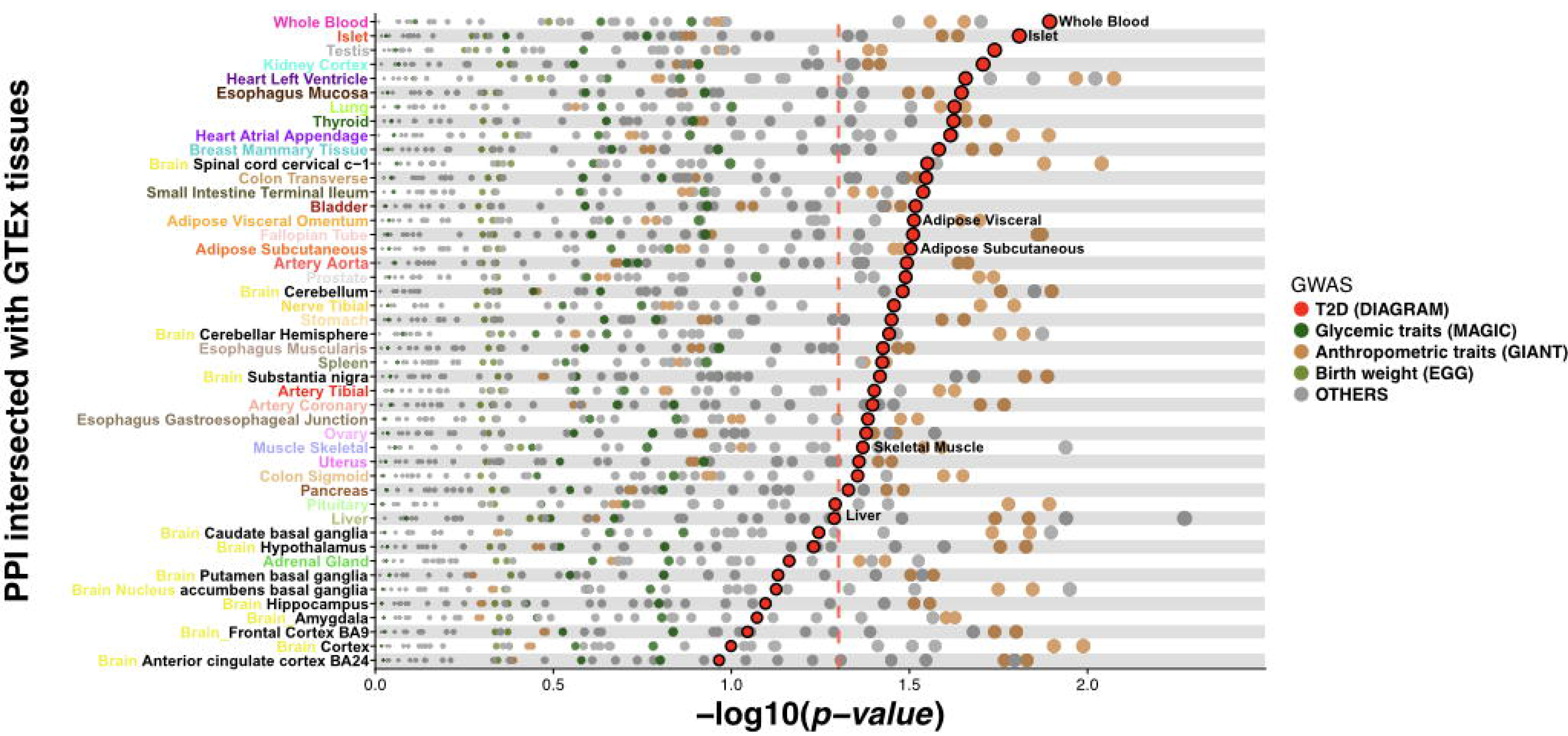
GWAS signal enrichment in tissue-specific interactomes. RNA-Seq data was used to filter the overall lnWeb3 network and generate in silico tissue-specific networks that maximise connectivity between GWAS interval genes. Linking nodes within these networks were then tested for enrichment for GWAS signals using a permutation scheme. Each dot in the figure depicts the ™log_10_ p-value for enrichment for signals in a given GWAS dataset, for each of the 46 tissues. Dot colors reflect the GWAS phenotypes with T2D in the larger red color. The dotted red line represents the nominal value of significance (*p*=0.05). Islet showed the second strongest enrichment signal for T2D.

### Further enhancement of model using GWAS locus subsets

To further refine the analysis, we took account of the multi-organ nature of T2D and, specifically, of evidence that it is possible, using patterns of association across T2D-related quantitative traits such as BMI, lipids and insulin levels, to define subsets of T2D GWAS loci which impact primarily on insulin secretion and those that perturb insulin action [47–49]. We reasoned that the former would be expected to show preferential enrichment within the islet-filtered PPI network. Accordingly, we built APCST networks (both generic and filtered for expression in islets exactly as above) formed from the sets of high-PCS seed genes mapping to each of seven T2D GWAS locus subsets defined in two recent publications [48], [49].

In both the islet-specific (**Fig 4**) and the generic network (**S7 Fig**), the strongest signals for GWAS enrichment were seen for loci in the three subsets (beta-cell [BC] in [48]; acute insulin response [AIR] and peak insulin response in [49]) comprised of T2D GWAS loci which influence T2D risk primarily through a detrimental effect on insulin secretion (**Fig 4**; **S6 Fig**). In particular, there was striking enrichment in the islet-specific PPI network for linking nodes in analyses of the BC (*p*=3.9×l0^−5^) and AIR (*p*=1.9×l0^−4^) T2D GWAS locus subsets.

**Figure 4.**
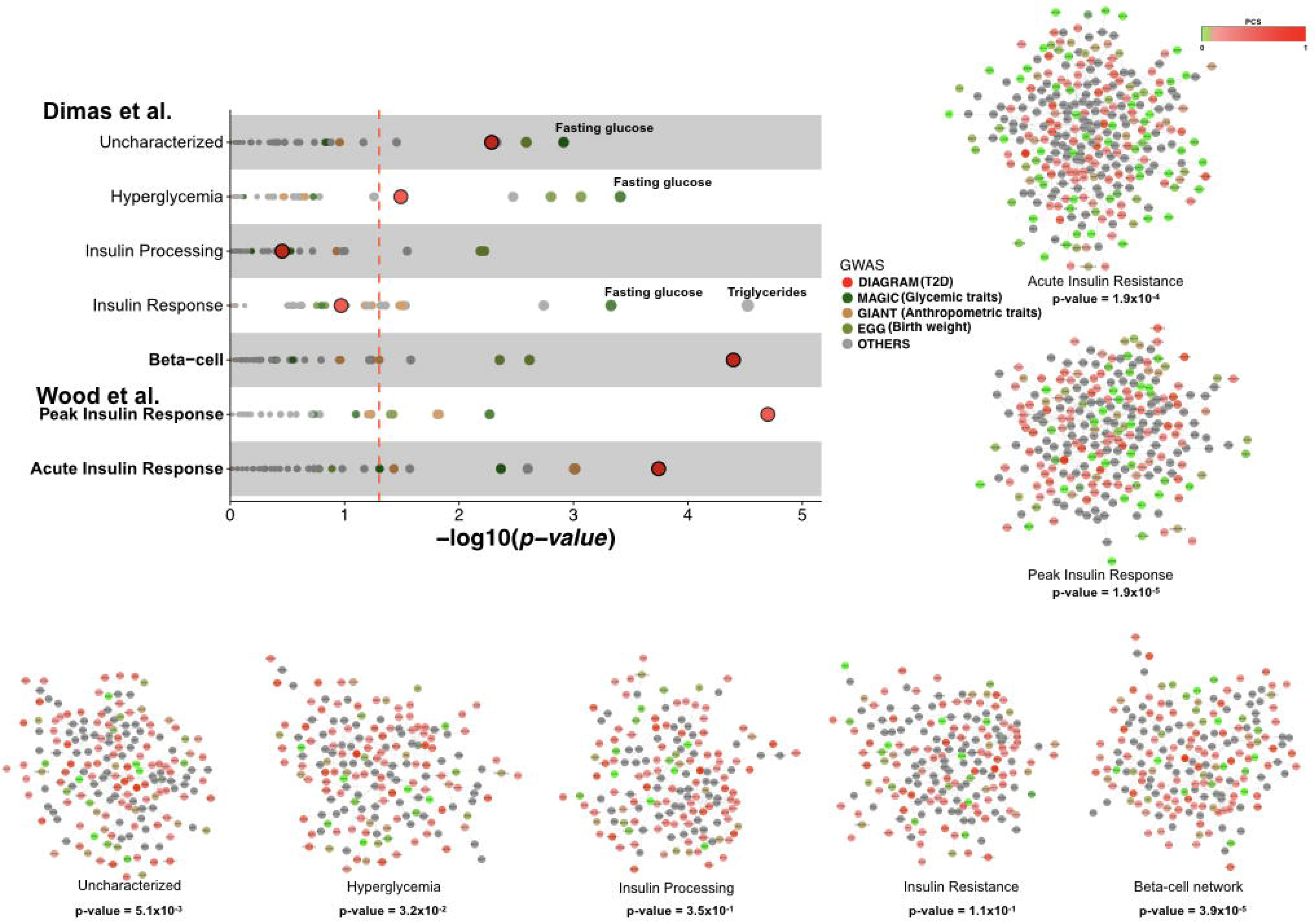
GWAS signal enrichment in islet-specific network derived from T2D GWAs subsets. We built APCST networks filtered for islet RN A-expression for each of the subsets of T2D GWAS loci defined by shared mechanistic mediation (refs[48], [49]. Enrichment in GWAS signals for linking nodes only was tested using a permutation scheme. Each dot in the figure depicts the ™log_10_ p-value of enrichment for association signals in a particular GWAS analysis. The results for T2D GWAS enrichment for the APCST networks built around the different T2D GWAS subsets are also represented (large red dots). The dotted red line represents nominal significance (*p*=0.05). The strongest enrichment for T2D GWAS data in islet-filtered PPI data is observed for subsets of loci acting through reduced insulin secretion. In the cluster hairballs for the seven T2D GWAS locus subset categories, nodes are coloured according to their PCS with grey nodes representing linking nodes.

As before, we were interested to see whether this marked convergence of PPI signal (as assessed by the enrichment of T2D association signals in linking nodes) was T2D-specific. We therefore repeated the enrichment analysis using GWAS data from 33 additional traits. For each trait, we took the APCST networks generated using the seven T2D locus subsets and assessed the “linking” nodes in those networks with respect to enrichment for respective gene-wise association p-values. We found broad levels of enrichment for association signals for T2D-related phenotypes including (quantitative) glycemic traits, lipid levels, anthropometric and cardiovascular traits, which are consistent with known GWAS signal overlap. However, we saw very limited enrichment for other (non-diabetes related) traits. Furthermore, the patterns of enrichment were consistent with underlying physiological expectation: GWAS enrichment for anthropometric and lipid phenotypes was most marked in the APCST networks generated from the insulin-resistant subset of T2D loci (category “insulin response” in [48]), whilst T2D remained the most enriched phenotype for the subsets related to insulin secretion (**Fig 4**).

These analyses demonstrated that parallel efforts to refine the phenotypic impact of T2D GWAS loci, and the tissue-specificity of the underlying PPI dataset used to generate the APCST network, resulted in progressive, biologically-appropriate, improvement of the enrichment signal observed at the “non-seed” proteins represented within the network.

### Biological insights

To better understand the biological function of the highly-enriched PPI network generated by the intersection of islet-specific expression, and the subset of T2D GWAS loci acting through reduced islet function (henceforth, the “islet network”), we performed a Gene Set enrichment analysis using GO and REACTOME terms (**S4 Table**). Captured pathways included well-known biological processes of “glucose homeostasis” (p =1.5×l0^−4^), “regulation of WNT signalling pathway” (p=8.9×l0^−3^), “response to insulin” (p=6.2×l0^−4^), and “pancreas development” (p=3.0×l0^−5^).

This islet network included many “seed” genes with a high T2D PCS score (**Fig 4**) including *SOX4* ([PCS=0.62] at the locus usually named for *CDKAL1*), and *ATXN7* ([PCS=0.57] at the locus named for *ADAMTS9*). Some of the loci (e.g. *TLE4, CAGE1* and *GCK*) are represented by a single “seed” because the PCS for the highest-ranking gene exceeded 0.70. At other loci, this islet network does not include the gene with the highest PCS score for the respective GWAS signal, but instead features an alternative gene from the same locus on the basis of its better connectivity within the network. Examples such as the gene *TBS* [PCS=0.21] at the *ZBED3* locus, and *THRB* [PCS=0.43] at the *UBE2E2* locus, demonstrate how the PPI data provides information additional to that used to derive the PCS.

In addition, several of the linking nodes introduced into this islet network through their PPI connections represent interesting candidates for a role in T2D pathogenesis. Cyclin-dependent kinase 2 (CDK2), for example, has been shown to influence beta-cell mass in a compensatory mechanism related to age and diet-induced stress, connecting beta-cell dysfunction and progressive beta-cell mass deterioration [50]; YHWAG is a member of the 14-3-3 family, known to be signalling hubs for beta-cell survival [51]; and disruption of SMAD4 drives islet hypertrophy [52].

## Discussion

In this study, we set out to overcome two challenges that have impeded efforts to synthesise the biological information that is captured in the growing number of association signals emerging from GWAS studies. In the case of type 2 diabetes, for example, there are now well over a hundred independent common variant signals [6, 21], but most of these map to regulatory sequence, and the molecular mechanisms whereby these, individually and/or collectively, contribute to differences in T2D predisposition remain largely unresolved. A key question, of direct relevance to the opportunities for translational use of this information, is the extent to which, as the number of loci expands, there will be “saturation” or “convergence” of the biological mechanisms through which they operate, or whether, on the contrary, the range of networks and pathways implicated will continue to proliferate.

The first challenge concerns the identification of the effector transcripts through which the T2D predisposition effects at each of the GWAS signals (most obviously those that are regulatory) are mediated. We approached this challenge by integrating, for each of the genes within each of the GWAS signals, two types of data, one based around the fine-mapping of the causal variant, and the use of cis-eQTL data (in the case of regulatory variants) or direct coding variant inference to highlight the most likely effectors, the other making use of diverse sources of biological information concerning the candidate effector genes and their protein products. Using this framework, we were able to assign candidacy scores to each regional gene, and then to deploy these scores as summaries of diverse sources of data that could be propagated into subsequent network analyses. We recognise that, given the sparse nature of the data used, not all such candidacy assignments will be accurate. However, these scores provide a principled and objective way of synthesising current knowledge, and the framework allows for iterative improvements in candidacy assignments as additional sources of relevant data become available. These are likely for example, to include further refinements in fine-mapping, additional links from associated variants to their effectors arising from chromatin conformation analyses, detection of rare coding variant signals through exome sequencing, and genome-wide screens of transcript function.

The second challenge relates to the objective evaluation of the extent to which the strongest positional candidates at these GWAS loci occupy overlapping biological space. Standard approaches to network analysis applied to GWAS data ‒ such as gene-set enrichment [32], or co-expression analyses [53] ‒ were not an option for this study since source data relevant to these had already been factored into the assessments of positional candidacy. Instead, we focused on the relationships between positional candidates as revealed by protein-protein interaction data, which we considered to be independent of the data in the earlier stages. We used the enrichment of T2D association signals in linking nodes (i.e. proteins included in the network which did not map to known GWAS loci) as our principal metric of network convergence.

This strategy uncovered a highly-interconnected network associated with T2D, which was built around proteins involved in processes such as autophagy, lipid transport, cell growth, and insulin receptor signalling pathways. We were able to show that this signal of enrichment was enhanced when we constrained the generic PPI network to reflect only genes expressed in pancreatic islets, and, concomitantly, limited the set of GWAS loci to those at which the T2D predisposition was mediated by defective islet function. These analyses reinforce the importance of the pancreatic islet as a critical tissue for the development of T2D, and highlight multiple proteins (both those that map within GWAS loci, and those that fall outside) that are represented within this core islet network. These findings provide compelling hypotheses that can be explored further through direct experimental study, and also highlight the need to generate tissue-specific protein-protein interaction data. They also provide evidence to support a convergence of the mechanisms mediating predisposition across diverse T2D association signals.

Finally, these analyses demonstrate a valuable approach for the interrogation of large-scale GWAS data to capture biologically-plausible disease-specific processes, one which can readily be applied to other complex diseases.

## Acknowledgments

We would like to acknowledge Dr. Heiko Horn and Dr Loukas Moutsianas for their valuable comments and useful discussion about methods during the elaboration of this manuscript.

## Supporting information

**SI Fig. Correlation between β values and PPI network size**

We tested different values of β to characterise the impact on network size and the percentage of seed genes represented in the network. The figure depicts the relationship between β values and both the number of nodes and the percentage of seed genes in the optimal solution generated with the Steiner tree approach. We selected as optimal a β value of 10 which produces a network of ~150 nodes which contains at least 25 % of the seed genes.

**S2 Fig. Specificity of linking nodes in the final network.**

We assessed the specificity of each node in the final network solution by running the algorithm 100 times with the same parameter settings, but with random input data. We define specificity in this context as the complement of the percentage with which a given linking node from the final network appears in runs generated from random input data. 80% of linking nodes have a specificity exceeding 0.75, indicating that these linking nodes do not simply reflect generic hubs in PPI space.

**S3 Fig. Distribution of PCS and correlation of semantic and risk variant link scores**. a)

Distribution of PCS values for the 1,895 candidate genes (left histogram) and for the 451 prioritised candidate genes (those that, for each locus contribute collectively to at least 70% of the total PCS) (right histogram); b) the number of genes per locus stratified in terms of PCS ranges for the 1,895 candidate genes (left boxplot), and the 451 prioritised candidate genes (right boxplot); c) the correlation between the semantic and risk variant link scores for the 1,895 positional candidates.

**S4 Fig. Summary of characteristics of PCS values.**

a) Distribution of gene number per locus for the 1,895 candidate genes (top histogram); and for the 451 prioritised candidate genes (bottom histogram); b) example of the distribution of PCS scores for the *TP53INP1* locus under “nearest-gene” selection (top figure), “all gene” selection (median figure), and under our prioritisation method (bottom figure); c) Scatter plot displaying the correlation between maximum PCS values for each locus under “nearest gene” our prioritisation approach; d) distribution of the maximum PCS per locus with our prioritisation strategy.

**S5 Fig. Enrichment of GWAS signals in the final PPI network.**

Using the generic PPI network generated from optimisation of seed node connectivity, we performed GWAS signal enrichment analyses (linking nodes only) for 33 GWAS datasets including T2D. Each point represents the −log_10_ p-value of the enrichment signal for the specified GWAS dataset. The strongest signal of enrichment was observed between the generic network and T2D (*p*=0.0014), with other significant associations for related phenotypes such as BMI. The dotted red line represents nominal significance (*p*=0.05).

**S6 Fig. Correlations between tissue-specific PPI networks.**

The composition of 46 tissue-specific networks generated by filtering the generic PPI network using tissue-specific RNA-Seq abundance data, was compared by applying a Jaccard similarity index to network nodes. Many tissue-specific networks showed high similarity with grouping by higher-level tissue of origin (e.g. brain, artery).

**S7 Fig. GWAS signal enrichment in the PPI-generic network derived from T2D GWAs subsets.**

We built APCST networks using the generic PPI network for each of the subsets of T2D GWAS loci defined by shared mechanistic mediation (refs[48], [49]). Enrichment in GWAS signals for linking nodes only was tested using a permutation scheme. Each point in the figure depicts the −log_10_ p-value of enrichment for association signals derived from a particular GWAS analysis. The results for T2D GWAS enrichment for APCST networks built around the different T2D GWAS subsets are also represented (large red dots). The dotted red line represents nominal significance (*p*=0.05). The strongest enrichment for T2D GWAS data in the generic PPI network is observed for subsets of loci acting through reduced insulin secretion. In the cluster hairballs for the seven T2D GWAS locus subset categories, nodes are coloured according to their PCS with grey nodes representing linking nodes. Comparison with Figure 4 (the equivalent figure generated using the islet-filtered PPI network) demonstrates increased levels of enrichment for the subset of T2D loci influencing insulin secretion when filtering for nodes reflecting pancreatic islet expression.

**S1 Table. 101 loci and candidate genes by loci used to calculate the Positional Candidacy Score (PCS).** Note: We developed a framework to score the candidacy of genes mapping to GWAS association signals which aggregated data from multiple sources. The information collected fell into two categories. First, we used regression-based approaches to link disease-associated variants to their likely effector transcripts, using a combination of variant-based annotations and expression QTL data (Link score). Second, we scored each of the genes in these GWAS regions for disease-relevant biological function (Semantic score). We combined the two measures to generate a “positional candidacy score” (PCS) for each gene. Cumulative: cumulative frequency of PCS for each loci. References: Bibliographic references describing the loci as associated to type II diabetes.

**S2 Table. DisGeNet results.** MeSH = Medical Subject Headings; DPI score = disease pleiotropic index; DSI score = disease specific index; GDA Score = Gene-Disease Association Score; El = Evidence Score.

**S3 Table. Gene Set Enrichment Analysis by community.** GOID: Gene Ontology ID; GOTerm: Gene Ontology Term; Note: Gene Set Enrichment (GSE) of networks and sub-networks was performed with ClueGO using GO terms and REACTOME gene sets. The enrichment results were considered significant when bonferroni adjusted p-value < 0.05 and at least 3% of the genes contained in the tested gene set is included in the network. Gene sets were also grouped using kappa score into functional groups to improve visualization of enriched pathways.

**S4 Table. Gene Set Enrichment Analysis in Beta-cell Islet-specific network.** GOID: Gene Ontology ID; GOTerm: Gene Ontology Term. Note: Gene Set Enrichment (GSE) of networks and sub-networkswas performed with ClueGO using GO terms and REACTOME gene sets. The enrichment results were considered significant when bonferroni adjusted p-value < 0.05 and at least 3% of the genes contained in the tested gene set is included in the network. Gene sets were also grouped using kappa score into functional groups to improve visualization of enriched pathways.

